# From Genes to Glands: Unraveling the Pivotal Influence of NtAGL66, an AGAMOUS-like Transcription Factor, on Glandular Trichome Development in *Nicotiana tabacum*

**DOI:** 10.1101/2025.04.27.650782

**Authors:** Alice Berhin, Gabriel Walckiers, Manon Peeters, Belkacem El Amraoui, Charles Hachez

## Abstract

Glandular trichomes are specialized epidermal structures that play an essential role in plant defense by synthesizing, storing, and secreting specialized metabolites.

This study investigates the function of *NtAGL66*, an AGAMOUS-like gene in *Nicotiana tabacum*, uncovering its role in the development of secretory heads in long glandular trichomes. Expression profiling reveals that *NtAGL66* is specifically expressed in the developing secretory glands. Functional analyses show that *NtAGL66* overexpression promotes the differentiation of the secretory structure, while CRISPR-Cas9-mediated knockout significantly reduces the capacity of trichomes to form functional secretory glands, highlighting its essential role in trichome specialization.

Transcriptomic (RNA-seq) and functional genomic (DAP-seq) analyses indicate that NtAGL66 regulates also secondary metabolic pathways and is likely involved in broader transcriptional networks, including floral development. Notably, this includes genes such as NtTOE1, previously shown to control both floral organogenesis and glandular trichome formation in tomato. Moreover, NtAGL66 directly regulates the transcription factor NtGL2 through promoter binding.

By identifying an AGAMOUS-like gene as a key regulator of secretory gland development, this study offers novel insights into the genetic mechanisms underlying glandular trichome differentiation and specialized metabolite biosynthesis in *Solanaceae*.

## Introduction

Plant trichomes are specialized epidermal structures found on aerial organs and can be classified into different types: unicellular or multicellular, and glandular or non-glandular.

Non-glandular trichomes primarily provide a physical defense by forming a protective barrier, while glandular trichomes contribute both physically and chemically by synthesizing and secreting specialized metabolites such as terpenoids, flavonoids, alkaloids, and phenylpropanoids (Fahn, 2000; Glas et al., 2012; Schuurink and Tissier, 2020). These metabolites play key roles in plant defense against herbivores and pathogens, stress adaptation, ecological interactions and have valuable applications in pharmaceuticals and agriculture (Markus Lange and Turner, 2013; Huchelmann et al., 2017). Glandular trichomes exhibit diverse morphologies and secretory capacities and are commonly classified as peltate or capitate types, each with varying degrees of metabolic specialization (Werker, 2000; Tissier et al., 2017). By enhancing stress resilience and chemical defense mechanisms, glandular trichomes play an essential role in plant survival and reproductive fitness (Hauser, 2014). Environmental factors such as light intensity, temperature, and water availability significantly influence trichome density and secreted metabolite composition (Gianfagna et al., 1992; Yan et al., 2017; Guan et al., 2022; Lv et al., 2022).

Despite their economic and ecological importance, the molecular mechanisms governing glandular trichome differentiation and function remain only partially understood, particularly in comparison to the well-characterized non-glandular trichomes of *Arabidopsis thaliana* (Szymanski et al., 2000; Balkunde et al., 2010). Glandular trichome development is a multifaceted process regulated by a complex interplay of transcriptional regulators, hormonal signals, and cell cycle control mechanisms (Yang and Ye, 2013). Many transcription factors regulate multicellular trichome development in Asterid plants, highlighting the complex genetic networks underlying their formation (Han et al., 2022).

In *Arabidopsis*, trichome fate determination is regulated by a MYB-bHLH-WD40 transcriptional complex, with GLABRA1 (GL1), GLABRA3 (GL3), ENHANCER OF GL3 (EGL3), and TRANSPARENT TESTA GLABRA1 (TTG1) playing essential roles (Oppenheimer et al., 1991; Walker et al., 1999; Pesch & Hülskamp, 2009).

However, in *Solanaceae* species such as *Nicotiana tabacum* and *Solanum lycopersicum*, glandular trichome formation follows a transcriptional pathway differing from the regulatory mechanisms controlling the formation of unicellular non-glandular trichomes (Glover et al., 1998; Payne et al., 1999; Perez-Rodriguez et al., 2005) and is regulated by different sets of transcription factors (Chang et al., 2024). Recent studies in tomato underscore the existence of a finely tuned regulatory landscape distinct from that of unicellular trichome development (Wu et al., 2023; Wu et al., 2024). Key transcription factors governing glandular trichome development include several major families. Among them, MYB transcription factors play a central role, including SlMX1 in *Solanum lycopersicum* (Ewas et al., 2016; Wu et al., 2023), NbMYB123 in *Nicotiana benthamiana* (Liu et al., 2018) and GLAND CELL REPRESSOR (*SlGCR1* and *SlGCR2*) (Chang et al., 2024). In addition to MYBs, other transcription factor families contribute to trichome development. C2H2 zinc-finger proteins (ZFPs), such as *SlHAIR, SlZFP6, SlZFP8/SlHAIR2* (Chang et al., 2018; Zheng et al., 2021), and *NbGIS* (*GLABROUS INFLORESCENCE STEM*) (Liu et al., 2018), have been implicated. Similarly, bHLH transcription factors, including *SlMYC1* (Xu et al., 2018), and homeodomain-leucine zipper (HD-ZIP) IV proteins, such as *SlCD2* (Nadakuduti et al., 2012), *SlWOOLLY*, and *NbWOOLLY* (Yang et al., 2011; Yang et al., 2015), are also key regulators. Further regulatory elements include AP2/ERF transcription factors, including *LEAFLESS (SlLFS)* and *SlTOE1B* (Wu et al., 2023; Chang et al., 2024), WUSCHEL-related homeobox (WOX) proteins, such as *SlWOX3* (Wu et al., 2023), and TCP transcription factors, like *BRANCHED2a (SlBRC2a)* (Wu et al., 2024). Beyond transcription factors, cell cycle regulators also influence trichome formation. E3 ubiquitin ligases, such as *SlMTR1/SlCYCB2* and *SlMTR2*, along with *NbCYCB2* and *NtCYCB2*, play crucial roles in coordinating cell division and differentiation, reinforcing the intricate regulatory mechanisms governing multicellular trichome architecture in *Solanaceae* (Gao et al., 2017; Wu et al., 2020; Wang et al., 2021; Wu et al., 2023).

MADS-box transcription factors are well known for their roles in floral organ identity, fruit development, and root architecture (Riechmann and Meyerowitz, 1997; Gramzow and Theissen, 2010). Among them, the AGAMOUS-like (AGL) subfamily has been primarily associated with reproductive organ specification and floral transition (Bowman et al., 1989; Pnueli et al., 1994; Dreni and Kater, 2014).

Floral development in *Arabidopsis thaliana* is tightly regulated by interactions among MADS-box transcription factors, which define organ identity and timing through the ABCDE model (Soltis et al., 2007; Guo et al., 2015). These transcription factors ensure proper development of sepals, petals, stamens, carpels, and ovules (Pelaz et al., 2001; Ditta et al., 2004). LEAFY (LFY) and SUPPRESSOR OF OVEREXPRESSION OF CONSTANS1 (SOC1) integrate environmental cues to regulate floral meristem identity and activation of key floral regulators (Lee and Lee, 2010; Jin et al., 2021). SHORT VEGETATIVE PHASE (SVP) acts as a repressor, delaying the transition to flowering by inhibiting SOC1 and FLOWERING LOCUS T (FT), thereby blocking LFY activation.

Interestingly, the genetic program governing leaf development remains active during flower formation, it is partially modified by the action of floral homeotic proteins to establish reproductive organ identity (Ó’Maoiléidigh et al., 2013). Studies in *A. thaliana* have shown that AGAMOUS (AG) directly represses key regulators of trichome initiation, including *GLABRA1 (AtGL1)* and *ZINC-FINGER PROTEIN8 (AtZFP8)*, while promoting the expression of trichome repressors like TRICHOMELESS1 (TCL1) and CAPRICE (CPC) (Ó’Maoiléidigh et al., 2013). A perturbation of AG function results in the ectopic formation of trichomes on carpels, suggesting that AG-mediated suppression of trichome development is an integral part of floral organ identity maintenance (Ó’Maoiléidigh et al., 2013).

However, no AGAMOUS-like gene has been directly linked to glandular trichome formation, leaving unanswered the question of whether MADS-box transcription factors play any role in glandular trichome development.

In this study, we identify *NtAGL66*, an AGAMOUS-like transcription factor, as a previously uncharacterized regulator of glandular trichome development in *Nicotiana tabacum*. Our findings provide the first direct evidence that a MADS-box gene contributes to the development of the secretory heads of long-stalked glandular trichomes. By demonstrating that an AGAMOUS-like MADS-box gene plays a previously unrecognized role in glandular trichome formation, our study provides novel insights into the molecular networks controlling epidermal differentiation and contributes to a deeper understanding of epidermal specialization in *Solanaceae*, a plant family including plants of agronomical importance.

## Results

### NtAGL66 expression is confined to the developing trichome head during glandular trichome development

*Nicotiana tabacum* is an allotetraploid species derived from a natural hybridization between *Nicotiana sylvestris* and *Nicotiana tomentosiformis*. Since *N. tabacum* conserved both genomes, we have most of the genes in two homeologs copies, one from each genome. *NtAGL66* (LOC107788518) closest homolog is *NtAGL104* (LOC107759308) with 96% identity. *NtAGL66* is identical to LOC104119783 from *N. tomentosiformis* while Nt*AGL104* has 96% identity with LOC104119783 from *N. sylvestris*.

A spatio-temporal analysis was conducted to understand where *NtAGL66 and NtAGL104* are expressed within the plant. The study of their expression level across a range of tissue types in *Nicotiana tabacum*, using quantitative RT-qPCR with primers amplifying both genes together, showed no variability between leaves of different stages and very low expression level compared to internal control genes (Figure S1A) suggesting an absence or an expression only in a few cells.

Analysis of the transcriptional reporter line *pAGL66::nlsGFP-GUS* revealed that *NtAGL66* promoter is only active in the early stages of trichome head development, beginning at the one-cell stage of gland development (Figure 1A). Interestingly, no activity is detected in the developing stalks of the trichomes nor in trichome initials. This activity is also observed in the cell below the gland during the early stage of gland cell division. This cell-type specific activation in early stages of gland development could suggest that *NtAGL66* plays a foundational role in initiating the development of the glandular heads of trichomes. As the trichome head develops, the promoter activity is not restricted to a single cell but extends throughout all the cells that will eventually form the glandular head structure. Once the glandular trichomes have matured, typically evidenced by their presence on young leaves, the activity of the *NtAGL66* promoter diminishes, with no signal detected in mature leaf trichomes. This *NtAGL66* activity across the trichome head cells indicates that *NtAGL66* could be important for the development and formation of the glandular trichome, likely coordinating the cellular processes necessary for full maturation, rather than being involved in the ongoing production of metabolites in mature secreting structures.

**Figure 1:**
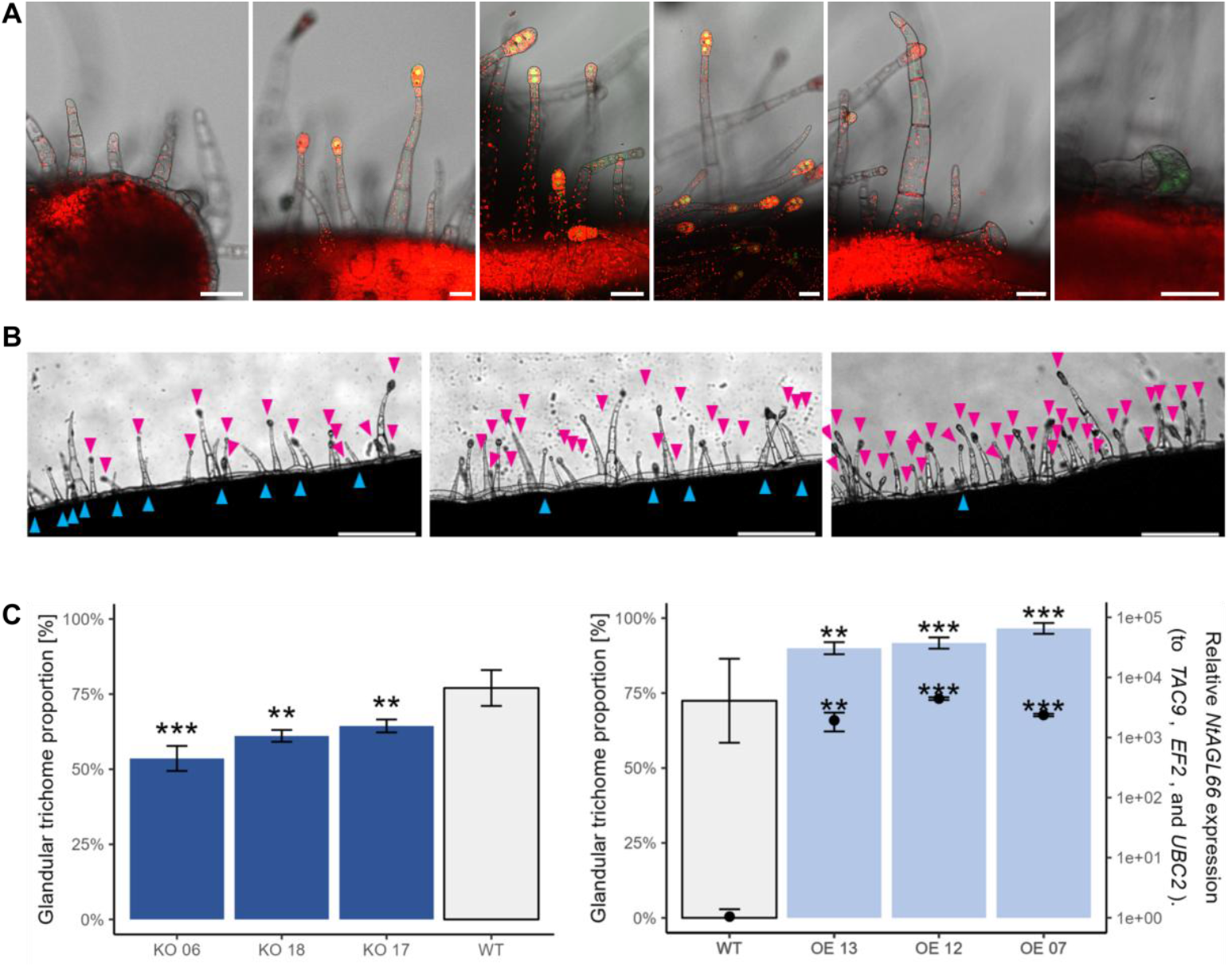
NtAGL66 is expressed in trichome head and influence the density of glandular trichomes. **(A)** Expression of AGL66 in the head of trichomes during their formation. Gene expression in transgenic plants expressing *pAGL66::nlsGFP-GUS* in the leaves of 6-week-old plants. From left to right: developing trichomes without head yet, trichomes with heads at different stages, non-glandular trichomes, small glandular trichomes. Scale bars is 50 μm. Yellow/Green, GFP; Red, Chlorophyll autofluorescence. **(B)** Pictures illustrating the trichome layout in WT, CRISPR-Cas9 mutants and overexpressing lines. Sale bars is 500 μm. Blue arrow head, non-glandular trichomes; Rose arrow head, glandular trichome **(C)** Proportion of glandular trichomes. Results are displayed as mean ± SD, n=3. Asterisks denote significant differences to WT condition as determined by Student’s t test: *p < 0.05, **p < 0.01, ***p < 0.001. Left graph includes as well with dots the expression level of NtAGL66 in the different overexpression lines. Gene expression levels are expressed relative to the WT and to the geometric mean of three different internal controls (*NtEF1a, NtTAC9, NtUBC2*). Results are displayed as mean ± SD, n=3. Asterisks denote significant differences to WT condition as determined by Student’s t test: *p < 0.05, **p < 0.01,***p < 0.001

### NtAGL66 influences glandular trichome head formation

*NtAGL66* and *NtAGL104* were edited using CRISPR-Cas9 (KO lines) and generated p*35S::AGL66* overexpression lines (OE lines). Modulating *NtAGL66* expression in transgenic plants revealed changes in the glandular vs. non-glandular trichome ratio, highlighting its regulatory role in glandular trichome formation.

While the ratio of glandular to non-glandular trichomes is around 75% in WT plants, this ratio drops to 50–65% in CRISPR-edited lines. In contrast, lines overexpressing *NtAGL66* show an opposite trend, with an increase up to 95%. (Figure 1B-C). The reduction of glandular trichomes in the absence of *NtAGL66* indicates that this gene is essential for their proper differentiation, while higher expression levels of *NtAGL66* enhance the formation of glandular heads. This suggests that *NtAGL66* plays an essential role in promoting the formation of glandular trichome heads.

Differential RNA-seq analysis on overexpressing and mutated lines resulted in high-quality mapping to the *N. tabacum* genome (Sierro et al., 2014) (Table S1), identifying respectively 1856 and 2052 differentially expressed genes (DEGs) (Fold change > 1.3 for the upregulated DEGs & Fold change<0.68 for the downregulated DEGs; p-value < 0.05) (Table S2).

Among the DEGs, 64 were shared between the 904 upregulated DEGs in the overexpressing lines and the 1006 downregulated DEGs in the knockout lines (Figure 2A). Out of those, we identified 10 transcription factors using the Plant Transcription Factor Database (Jin et al., 2017) (Figure 2D).

**Figure 2:**
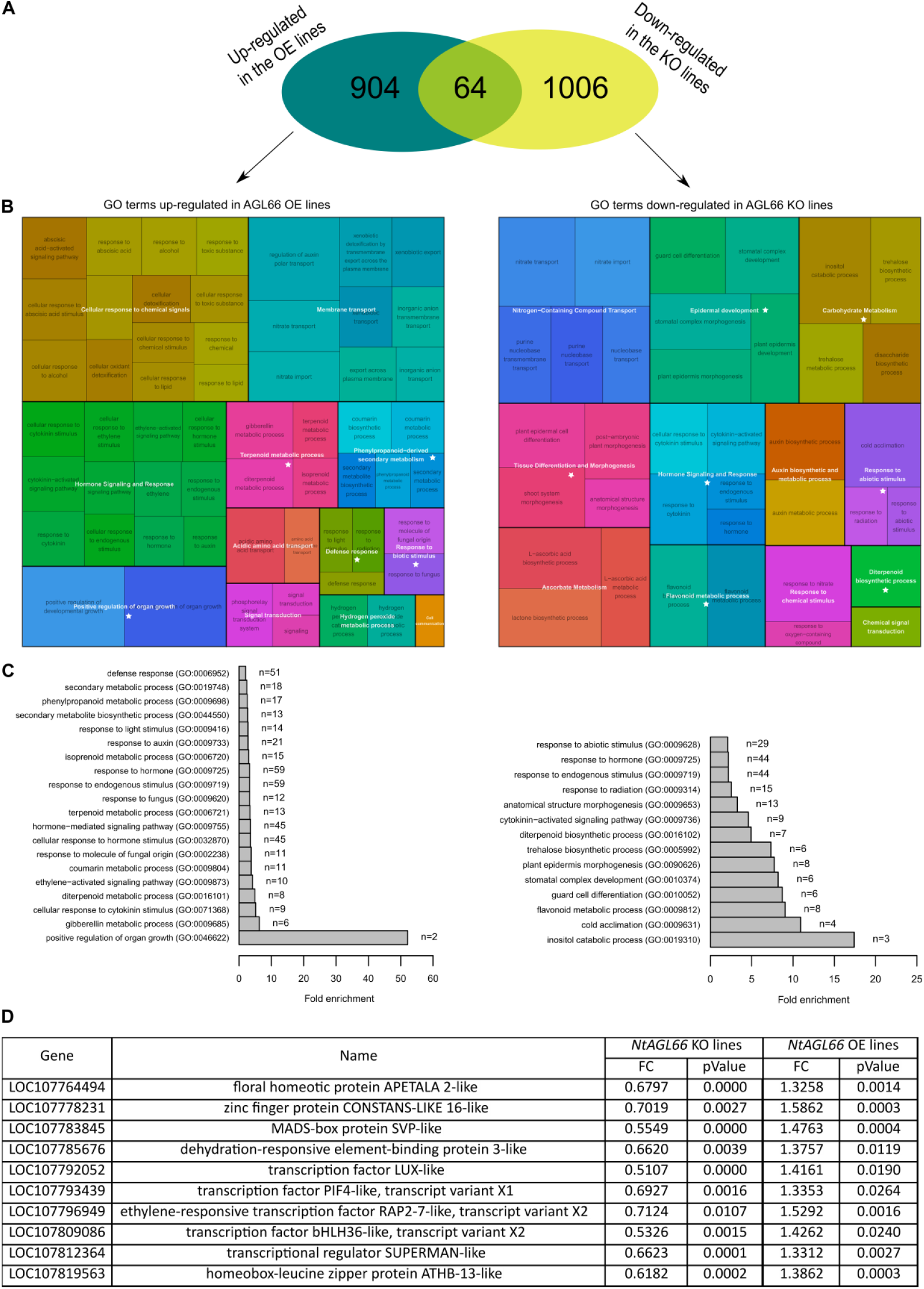
*NtAGL66* influence GO term categories linked to tissue development and metabolism of trichome secondary metabolites. **(A)** Overlap between the genes upregulated in the *NtAGL66* overexpression lines and in the mutated lines in comparison to WT. **(B)** Overview of the enriched categories using Rrvigo (Sayols, 2023), a tool to help visualize enriched GO terms. **(C)** Fold enrichment of the GO terms related to trichomes, selected based on the categories of the Rrvigo, annotated with a star in (B), redundant GO term was regrouped. (D) Out of the 64 genes commonly expressed between the genes upregulated in the NtAGL66 overexpression lines and in the mutated lines, 10 were transcription factors, here is the list.

Several of these transcription factors belong to families that have been previously characterized as involved in trichome development, including the AP2/EREBP ethylene-responsive element binding family, bHLH family, C2H2 zinc finger family and HD-ZIP family. Interestingly, the *RAP2-7-like* gene (LOC107796949; Figure 2D), that we named *NtTOE1*, is the closest ortholog is *SlTOE1B* in tomato (Chang et al., 2024), known to encode a positive regulator of glandular cell development. In *tomato*, high expression of the *GLAND CELL REPRESSOR (SlGCR1/2)* inhibits glandular head formation, leading to the development of non-glandular trichomes. SlTOE1 forms a complex with SlGCR1/2, relieving this inhibition and promoting gland formation. SlGCR1/2 inhibits trichome head formation by repressing the expression of *LEAFLESS (SlLFS)*, gene coding for an AP2-type transcription factor first identified as a regulator of peltate trichome fate (Wu et al., 2023). *NtLFS* is also overexpressed in our data following overexpression of *NtALG66* (Figure 2 and Table S2). Increased *NtAGL66* expression leads to upregulation of *NtTOE1*, which suppresses the activity of a potential *NtGCR* repressor, thereby promoting *NtLFS* expression and facilitating glandular trichome formation. The closest homologs to *SlGCR1/2* in tobacco were unchanged.

several transcription factors with potential roles in floral development and differentiation were identified as well. From the ABCDE model, we observed that the A-function gene *NtAP2* is directly correlated with Nt*AGL66* expression, whereas *NtAP1-like* is inversely correlated, similar to its two redundant genes *NtCAL* and *NtAGL8* (Ferrándiz et al., 2000). The E-function gene *NtSEP1* and *NtSEP4* is also inversely correlated. Additionally, key regulators such as *NtLEAFY* and *NtSOC1* are inversely correlated, whereas *NtSVP* is positively correlated.

It is also interesting to notice that cis-abienol synthase (LOC107827720 and LOC107830721) and squalene synthase (LOC107760995) are increased when there is more trichomes suggesting the functionality of the new forming head.

Among the 1046 upregulated DEGs in the knockout line and 952 downregulated DEGs in the overexpression line, 91 genes were common (Table S2, Figure S2). Among them, 27 are transcription factors, including 15 MADS-box genes. This suggests that the change in *NtAGL66* expression is compensated by other MADS-box transcription factors, as we observe a decrease in their expression in the OE lines and an increase in the KO lines. This compensation/coregulation could explain the relatively weak KO phenotype.

In GO term enrichment (Panther) (Figure 2B-C, Table S3), we observed that genes upregulated in the OE lines and downregulated in the KO lines were associated with tissue development including respectively for example GO term like plant epidermis morphogenesis (GO:0090626) and positive regulation of organ growth (GO:0046622), highlighting a role of *NtAGL66* in developmental processes, potentially including glandular head formation (Figure 2B-C). Additionally, were observed enriched GO terms related to the metabolism of trichome secondary metabolites such as terpene, flavonoid, alkaloids and carbohydrate, further supporting its involvement in establishing a secretory structure. We also identified enriched GO categories linked to responses to biotic and abiotic stresses, highlighting the functional role of trichomes in plant defense (Figure 2B-C).

### NtAGL66 regulates trichome development through NtGL2

DNA Affinity Purification Sequencing (DAP-seq) was performed to identify the direct transcriptional targets of *NtAGL66* in *N. tabacum* (Bartlett et al., 2017). This analysis allows the mapping of DNA-protein interactions by capturing genomic DNA fragments that are bound by a specific protein of interest, in this case the NtAGL66 protein, providing insights into the downstream gene network regulated by *NtAGL66* and its role in a complex regulatory network. Analysis of the binding site locations surrounding annotated genes revealed that, a significant majority (79.3%) of the interactions were localized to regions upstream of the START codon, encompassing promoter regions and 5’ untranslated regions (UTRs), and to introns (Figure 3A).

**Figure 3:**
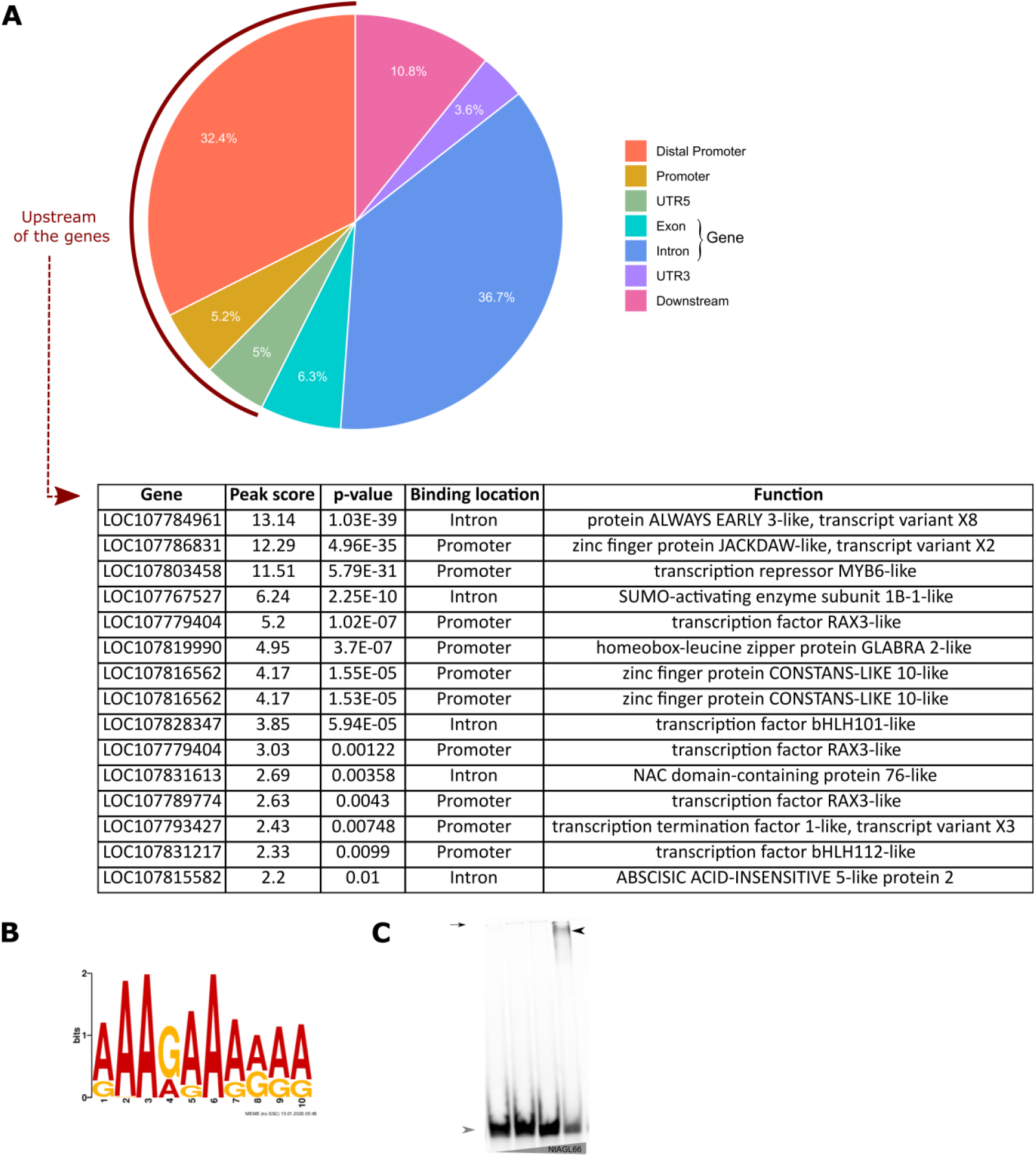
Identification of the direct targets of NtAGL66 using DAP-seq. **(A)** Distribution of NtAGL66-binding peaks relative to gene structures. Distal promoter, −3kb up the −0.5kb before the START codon; promoter, 0.5 kb up the UTR5; downstream UTR3 to 1kb after the STOP codon. **(B)** Top 13 list of the most significantly enriched region corresponding transcription factors which are bound by NtAGL66 in the region before the START codon or in introns. **(C)** The top-scoring motif, AAAGAAARAA was identified with an e-value of 1×10^−28^, according to MEME analysis. Gel shift assay shows a binding of NtAGL66 to the fluorescent DNA substrate: AAAGAAAAAA; Protein concentration were used: 0, 0.06, 0.6 and 11.92 μg of protein. black arrow head, bound DNA; Grey arrow head, unbound DNA; Small black arrow, wells.

Reads were filtered and mapped to the *N*.*tabacum* genome with an average mapping of 98% suggesting a high quality of the samples (Sierro et al., 2014) (Table S4). Hits were identified by comparing against the negative control (nlsGFP protein), using criteria of a peak shape score greater than 2 and a p-value < 0.01, to delineate significant binding regions (Table S5).

Among all the regions significantly enriched in the analysis, 12,46 % (444 hits) were in regions surrounding annotated genes (3kb before the start codon and 1kb after the stop codon respectively) corresponding to 376 distinct genes. The rest are in intergenic regions, that could still include unannotated genes, and were not further included in the subsequent analysis.

Direct targets of NtAGL66 bound in the intron or the promoter region highlighted the involvement of 12 transcription factor families in the NtAGL66-regulated pathway, including Zinc Fingers, MYBs, HD-ZIPs and bHLH (Figure 3A, Figure S3). These families are known to play essential roles in trichome development across different plant species. This suggests that NtAGL66 likely orchestrates the precise spatial and temporal expression of genes necessary for the formation and proper differentiation of secreting glandular trichome heads.

Our findings indicate that NtAGL66 regulates trichome development by directly targeting NtGLABRA2 (LOC107819990), ortholog of AtGLABRA2, SlGL2 and CsGL2, well-documented regulators of trichome initiation in *Arabidopsis*, tomato and cucumber plants respectively (Rerie et al., 1994; Yang et al., 2011; Liu et al., 2016), indicating that it might control the expression of NtGL2 and indirect control the development of the trichome (Figure 3A).

Analysis of *NtGL2* expression using the reporter line (*pGL2::nlsGFP-GUS*) revealed promoter activity in the developing trichome stalk but not in the gland, indicating that *NtGL2* and *NtAGL66* do not have overlapping expression patterns and are likely not acting synergistically (Figure 4).

**Figure 4:**
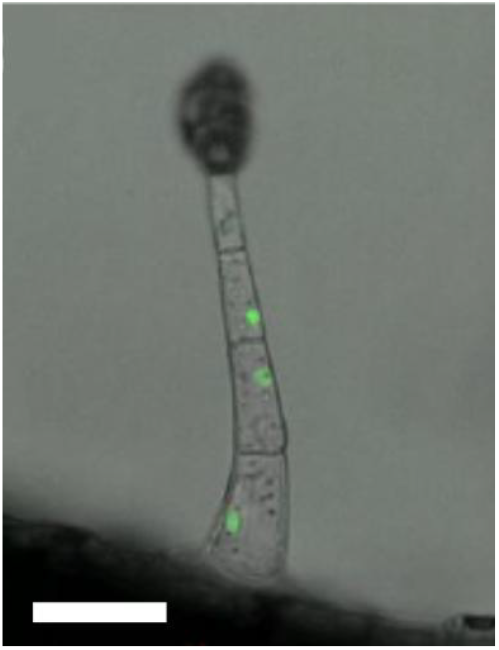
*NtGL2* is expressed in the trichome stalk. Expression of *NtGL2* in glandular trichomes during their formation. Gene expression in transgenic plants expressing *pGL2::nlsGFP-GUS* in the leaves of 6-week-old plants. Scale bars is 50 μm. Green, GFP.

Using the MEME suite for motif discovery (Bailey et al., 2015), we identified specific DNA sequences preferentially bound by NtAGL66: AAAGAAARAA motif, with an e-value of 1×10^−28^ across 215 binding sites (Figure 3B). Motif binding sites were confirmed using electrophoretic mobility shift assay (EMSA) confirming a sequence-specific binding of NtAGL66 to the AAAGAAAAAA motif (Figure 3C), this motif is fairly similar to known MADS binding motifs, the CArG box (CC(A/T)7GG) and variants (Folter and Angenent, 2006; Aerts et al., 2018).

## Discussion

The regulatory mechanisms underlying the development of glandular trichomes in *Nicotiana tabacum* represent an essential area of research due to their significant roles in plant defense and secondary metabolite production. Our comprehensive study sheds light on the role of the AGAMOUS-like gene, *NtAGL66*, in orchestrating the specialization of these structures.

### NtAGL66 has a role in the formation of trichome glandular head

The expression of *NtAGL66* displayed a temporal and spatial specificity, being confined to the early stages of glandular trichome head development (Figure 1A and S1). This expression pattern indicates that NtAGL66 plays a role in the differentiation of glandular trichome heads. Additionally, the decline in promoter activity in mature glandular trichomes highlights its specific function in their development, rather than in the later stages of metabolite production.

Reverse genetics employing CRISPR-Cas9 genome editing and overexpression strategies further confirmed *NtAGL66* regulatory function. Alterations in *NtAGL66* expression impacted the ratio of glandular to non-glandular trichomes, reinforcing its role in glandular trichome head formation (Figure 1B-C).

In tomato, glandular cell development is regulated by the positive factor SlTOE1B, which forms a complex with SlGCR1/2 to relieve repression of *SlLFS* (Wu et al., 2023; Chang et al., 2024). In our RNA-seq data (Figure 2, Table S1-S2), we identified *NtTOE1*, a homolog of *SlTOE1B*, suggesting a mechanistic link between *NtAGL66* and glandular trichome formation in *Nicotiana tabacum*. Overexpression of *NtAGL66* led to increased *NtTOE1* and *NtLFS* expression, supporting a model in which *NtAGL66* promotes glandular trichome development by relieving a potential NtGCR-mediated repression of *NtLFS* via NtTOE1.

Among the NtAGL66 direct transcriptional targets (Figure 3, Table S4-S5), *NtGLABRA2* (LOC107819990) stands out as a key candidate due to its homology with *AtGLABRA2, SlGL2*, and *CsGL2*, well-characterized regulators of trichome initiation in *Arabidopsis*, tomato, and cucumber, respectively (Rerie et al., 1994; Yang et al., 2011; Liu et al., 2016). With *NtAGL66* expression in the gland of developing trichome and the expression of *NtGL2* in the trichome stalk but not in the gland (Figure 4), we hypothesize that NtAGL66 act as a repressor of the expression *NtGL2* in the tip of the trichome leading the formation of a gland instead continuing the trichome growth division (Figure 3A).

Our results establish NtAGL66 as a key regulator of glandular trichome head formation, acting through multiple transcription factors, including NtTOE1 and NtGL2. Additionally, the identification of NtGL2 as a direct target suggests a conserved regulatory mechanism across species.

### NtAGL66 is a negative regulator of genes involved in flower development

AtTOEs are also known to act as key regulators that integrate developmental and environmental cues to control flowering in *Arabidopsis* (Zhang et al., 2015). *NtTOE1* is not the only flowering-related gene identified in our RNA-seq as several other transcription factors with potential roles in floral development and differentiation were detected (Table S2).

NtAGL66 activity modulated the expression of tobacco orthologues of such genes characterized primary as related to flowering (Table S2). It appears that *NtAGL66* expression correlates with *NtSVP*, which could inhibit *NtSOC1* activation of *NtLEAFY, NtAPETALA1 (NtAP1)* and *NtCAULIFLOWER* similarly as in *Arabidopsis* where *AtSVP* represses *AtAP1, AtPISTILLATA*, and *AtAPETALA3* expression (Meng et al., 2025). Similarly, *NtAP1* is inversely correlated with *NtSVP* and *NtAGL66* expression, while *NtPI, NtAP3*, and *NtFT* were downregulated when *NtSVP* was upregulated and *NtAGL66* was knocked out. *NtSOC1* could regulate *NtAGL42*, which promotes flowering in the shoot apical and axillary meristems (Dorca-Fornell et al., 2011), both inversely correlate with *NtAGL66* expression.

Additionally, genes positively correlated with *NtAGL66* expression include *ATHB13*, a homeodomain-leucine zipper I (HD-ZIP I) transcription factor whose ortholog in citrus has been shown to negatively regulate flowering (Ma et al., 2020), and *SUPERMAN* (*SUP*), a transcription factor known whose Arabidopsis orthologs regulates leaf cell division and expansion in tobacco (Bereterbide et al., 2001) and controls flower development and cell division in *Arabidopsis* (Hiratsu et al., 2002).

This suggests that NtAGL66 negatively regulates floral development genes, likely by upregulating NtSVP, a known repressor of flowering. Matias-Hernandez et al. (2016) showed that key flowering genes not only influence the transition to reproductive development but also modulate trichome initiation on rosette leaves, while trichome activators can reciprocally affect floral induction. Indeed, flowering and unicellular trichome development share transcriptional regulators: typical genes linked to trichome development (e.g., AtGL1, AtGL3, AtGIS, AtGIS2, AtZFP8, AtCPC) also influence floral transition, while MADS-box genes (AtFT, AtSOC1, AtFLC) regulate both processes (Wada and Tominaga-Wada, 2015; Matias-Hernandez et al., 2016). This crosstalk underlines a relationship between trichome specialization and floral induction, underscoring conserved genetic network that integrates hormonal and developmental signals to coordinate multiple plant developmental processes (Matias-Hernandez et al., 2016).

### NtAGL66 relates to trichome secondary metabolites production and export

Our findings suggest that *NtAGL66* not only governs glandular trichome differentiation but also contributes to their functional specialization in secondary metabolite biosynthesis.

The increased expression of cis-abienol synthase and squalene synthase in *NtAGL66* overexpressing plants with higher trichome density indicates that the newly formed glandular heads are actively engaged in terpenoid metabolite production (Table S2). Enriched GO terms included tissue development and metabolism of secondary metabolites such as terpenoids, flavonoids, and alkaloids, all of which are integral to trichome function and plant defense (Figure 2B-C).

Further insights into NtAGL66-mediated regulation come from the overlap between RNA-seq and DAP-seq datasets, which, although limited, highlight key downstream targets. Notably, two downregulated genes in the KO lines that were also bound by *NtAGL66* in the DAP-seq are of particular interest: *UDP-GLYCOSYLTRANSFERASE 71K1-LIKE* (*LOC107796811*) and *NON-SPECIFIC LIPID-TRANSFER PROTEIN 1* (*LOC107812329*). UDP-glycosyltransferases (UGTs) play an essential role in modifying secondary metabolites, including flavonoids and terpenoids, with glycosylation being an essential step in terpenoid biosynthesis within glandular trichomes (Lu et al., 2023). Similarly, non-specific lipid transfer proteins (nsLTPs) are associated with cuticle formation, lipid transport, and secretion of specialized metabolites. *NtLTP1*, a close homolog, is expressed in long glandular trichomes and contributes to lipid secretion outside of trichomes (Choi et al., 2012; Pottier et al., 2020). These findings suggest that *NtAGL66* regulates both the glycosylation and export of secondary metabolites in trichomes.

These results collectively support a model in which *NtAGL66* not only promotes glandular trichome formation but also orchestrates their biochemical specialization by regulating genes involved in secondary metabolite biosynthesis and transport. Future studies focusing on the biochemical activity of *NtAGL66*-regulated enzymes and their impact on metabolite accumulation will further elucidate the functional significance of this regulatory network.

## Conclusion

Our study highlights the important role of NtAGL66 in glandular trichome in *Nicotiana tabacum*. NtAGL66 emerges as a key regulator of secretory cell differentiation in developing glandular trichomes, acting within a complex genetic network. These findings lay the groundwork for future research aimed at leveraging glandular trichomes to enhance plant resilience and boost specialized metabolite production.

## Acknowledgments

We are grateful to Aubry Fenestre for the generation of the reporter lines of *NtGL2* promoter.

## Author Contributions

AB conducted the foundational experiments for this study and drafted the manuscript. GW performed trichome counts and EMSA and MP performed GC-MS experiment. BEA offered essential support in analyzing the RNA-seq and DAP-seq data. CH conceived and supervised the whole project and was actively involved writing of the manuscript.

## Data availability

RNA-seq and DAP-seq data sets are available upon request.

## Material and Method

### Generation of constructs

The *NtAGL66* (LOC107788518) coding sequence was amplified by PCR from cDNA (Table S6), and recombined into a pDONR221 vector to create the pENTRY L1-NtAGL66-L2. p35S::NtAGL66 was produced by LR Gateway reaction recombining the corresponding entry clone with p35S promoter into the pH7m34GW (Karimi et al., 2002).

To generate pENTRY L4-pNtAGL66-R1, 2401 kb fragment upstream of the NtAGL66 coding sequence amplied from genomic DNA (Table S6) and cloned into pDONR L4-L1r using KpnI and XbaI restriction site. pNtAGL66::nlsGFP-GUS was generated by recombining pENTRY L4-pNtAGL66-R1 with pENTRY L1-nlsGFPGUS-L2 (Berhin et al., 2019) into the pH7m34GW (Karimi et al., 2002).

For the CRISPR-Cas9, two guide RNAs (Table S6) were chosen to target exonic regions of NtAGL66 and NtAGL104. They were ordered as a gblock (GeneScript) in way that they were separated with a scaffold and both preceded by a tRNA. It was cloned by restriction digest and ligation to from P2P3r-AtU6-tRNA-gRNA-scaffold-tRNA-gRNA-scaffold. This plasmid, L4-pCYCD3-L1 and pDONR221-CAS9-tagRFP-T35S were transferred to the pH7m34GW plasmid via an LR Gateway reaction (Wang et al., 2020).

For the protein expression needed in the DAP-seq, pENTRY L1-NtAGL66-L2 was generated here above and pENTRY L1-nlsGFP-L2 (Karimi et al., 2002) were transferred through a LR Gateway reaction into its final pIX-HALO plasmid (Bartlett et al., 2017) to form pIX::HALO-NtAGL66 pIX::HALO-nlsGFP. For recombinant protein expression for the EMSA, the coding sequence of NtAGL66 was amplified (Table S6). The PCR amplicon and the pQE60 plasmid (Qiagen) were digested with BglII and NcoI-HF restriction enzymes and ligated together.

### Plant Material and growth conditions

*N. tabacum* cv Petit Havana (SR1) was used for stable plant transformation. For *in vitro* culture seeds were surface-sterilized with chlorine gas and sown on half-strength Murashige and Skoog agar plates (4.4 g/L MS basal medium (MP biochemicals, 92610024), 30 g/L sucrose, 8 g/L agar, pH 5.6 (KOH)). After 3 days at 4°C for stratification, the plates were transferred to a growth chamber with a 16-h-light/8-h-dark regime, 35 μmol photon/m^2^s and a temperature of 25±2^°^C.

For plant propagation and phenotyping purposes, seedlings were either obtained from *in vitro* cultures and placed in Jiffy for a week or were germinated directly in Jiffy for two weeks, before being transferred to pots containing potting soil and placed in a phytotron (25 °C, 16 h photoperiod, 200 μmol photon/m^2^s).

### Generation of stable transgenic lines

All constructs in the destination vectors were introduced into *Agrobacterium tumefaciens* strain GV3101. Constructs were transformed into tobacco using a modified version of the *Agrobacterium tumefaciens*-mediated transformation of leaf disks to generate transformants from (Horsch et al., 1989). 0.5 cm^2^ pieces of leaves were cut from *in vitro* grown WT plant and injured using a syringe needle previously immersed in the *agrobacterium* solution (*Agrobacterium* liquid culture centrifuged at 6000g for 2min and resuspended in 1ml of 10mM MgSO4 three times up to the last time where cells were resuspended in 300µL of MgSO4 and 1,5 µL of 200 mM acetoseringone (Sigma, D134406). Four days after, the infected leaf pieces were washed in three successive bath of a solution containing MS liquid medium with 500mg/L cefotaxin (Phytotech, C380), 400mg/L ampicillin (Roth, K029) and plant selection antibiotic (100mg/L of kanamycin (Roth, T832.2) or 30mg/l of hygromycin (Duchefa, H0192.0001)) and placed during minimum three weeks on solid plates containing 1/2MS medium with the same antibiotics, 0,2 mg/L 3-indol-acetic-acid (IAA) (stock in ethanol, Sigma, I2886-5G) and 2,2 mg/L 6-benzylaminopurin (BAP) (stock in DMSO, Sigma, B3408-1G) in order to promote the callus generation. Once the calli were well formed, they were cut and placed on solid plates containing MS medium with 250mg/L cefotaxin, 400mg/L ampicillin, plant selection antibiotic (15mg/L hygromycin) and 0.2 mg/L BAP in order to promote leaf generation. After three weeks, seedlings appearing on the callus, were cut off and transfer on solid plates containing MS medium with the previous antibiotic and only 0.2 mg/L IAA in order to promote the roots generation. After the development of the roots, seedlings were transferred to Jifi and then to soil. Lines generated for this work were plants carrying pNtAGL66::nlsGFP-GUS, pNtGL2::nlsGFP-GUS, p35S::NtAGL66, NtAGL66 CRISPR-Cas9 constructs.

T1 seeds from p35S::NtAGL66 and CRISPR *agl66* were already screened for glandular trichome phenotype. Overexpressing lines were tested for the expression level of *NtAGL66* (Table S6). The mutation of the CRISPR lines were tested by amplifying the gene (Table S6), sequencing the zone around the gRNAs and analyzing the mutation using Tide (Brinkman et al., 2014)

### RNA-seq sample preparation

T1 plants from p35S::NtAGL66 and CRISPR *NtAGL66* lines were confirmed for genotype and phenotype. The most interesting line of each was selected for preparation of samples for RNA-seq. They were grown first on Jiffy for two weeks then in big pots for 4 weeks. The samples were harvested on 6 weeks old (+-25cm in height) and were composed of all leaves of max 2 cm.

### Total RNA isolation

For the RNA extraction, the collected tissues were frozen in liquid nitrogen directly after harvesting.

For the RNAseq, smallest leaves (up to 2cm) were harvested per sample (3 replicates).

For expression study of NtAGL66, different tissues including leaf primordia (all leaves <1.5 cm), young leaves (one leaf around 10cm), mature leaves (one leaf around 20cm), mature trichomes (harvested by flash-freezing and brushing of a mature leaf) of 6-week-old plants (+-25cm height). For each of the harvested tissues, tissues from three plants were mixed to form a sample and four samples were harvested to have four biological replicates.

Samples were grinded using a mixer mil machine (Retsch GmbH) for 1 min at maximum speed (30 hertz). RNA was extracted with the Spectrum Plant Total RNA Kit (Sigma, STRN250-1KT) with on-Column DNase I Digestion set (Sigma, DNASE70-1SET).

### Gene expression level

1 µg ARN were reverse transcribed using M-MLV Reverse Transcriptase (Promega; M170B) and oligo(dT)18) following the manufacturer’s instructions. Real-time quantitative PCR analyses were then performed on the StepOne Real Time PCR Systems (ThermoFisher Scientific). qPCR reaction mix were prepared using the qPCR Master MIX Plus for SYBR Assay ROX (Eurogentec, RT-SN2X-06+). Amplification conditions included an enzyme activation step of 15 min at 95°C, followed by 35 cycles of 30 sec at 95°C and 1 min at 60°C, and a final 1 min at 95°C step. Dissociation curve analysis was done from 65°C to 95°C for 10 sec. Normalization was done by geometric averaging of multiple internal control genes: NtEF2α, NtACTIN, NtUBC2 (Vandesompele et al., 2002). The relative expression levels were calculated based on the 2^−ΔΔCT^ (Schmittgen and Livak, 2008). Primer sequences used are listed in Table S6.

### NGS sequencing and data processing for the RNA-seq

The RNA-Seq libraries were generated and sequenced by Genewiz (Azenta life science) using Illumina NovaSeq sequencing plateform. A minimum of 158 million 150 bp paired-end reads were obtained per sample (Table S1). Using CLC Genomic Workbench (Qiagen), reads were cleaned and trimmed of adaptors, ambiguous nucleotides, homopolymers in 3’ and 5’ and for quality (PHRED30) and minimum length. They were then mapped to the *Nicotiana tabacum* TN90 reference genome (GCF_000715135.1_Ntab-TN90_genomic) (Sierro et al., 2014). Identification of differentially expressed genes (DEGs) was based on normalized gene expression calculated as counts per million mapped reads (CPM), analyzed with a log2 fold change (FC) >1 or <-1 and a p-adj<0.05 using Rstudio with DESeq2 package (Love et al., 2014).

The resulting DEGs were uploaded to PANTHER classification system to look for significantly enriched GO (Gene Ontology) terms in the dataset (Mi et al., 2019; Thomas et al., 2022).

### DAP-seq sample preparation

Leaf primordia harvested from three independent WT plants were flash-frozen, grinded and DNA was extracted using the Wizard Genomic DNA Purification Kit (Promega; A1120). The DAPseq experiment was executed following the protocol from Bartlett et al. (2017) with some small modifications. Briefly, DNA was fragmented using a sonicating bath for 20 min (Elam S10H Elmasonic) and fragment size was verified on gel. Then the DNA fragments was subject to end-repair (Fast DNA end repair kit, Thermoscientific, K0771), A-tail reaction using Klenow enzyme (NEB, M0212S), ligation to the adaptor (Promega, M1794) and DNA concentration measurement (Qubit dsDNA Assay kit, Qiagen, Q32850). HALO-NtAGL66 and HALO-nlsGFP (negative control) were expressed independently three times using the TnT® Coupled Wheat Germ Extract System (Promega, L41030). Expression was confirmed by Western-blot using Monoclonal Antibody Raised Against the HaloTag® Protein (Promega, G9211) and Goat anti-mouse IgG (H+L) HRP conjugated (Chemicon, AP308P). Proteins were bound to the 20μl of Magne® HaloTag Beads (Promega, G7882) per sample, washed and put in contact with 1000 ng of DNA for 1h at 25°C. Each replicate of proteins was put in contact with one replicate of DNA. Beads were washed and DNA was eluted in EB at 98°C. Index and adaptor were added to the DNA fragments using Phusion® High-Fidelity DNA polymerase (NEB, M0530L) and NEBNext® Multiplex Oligos for Illumina® primers (NEB, E6440S), and DNA check on gel and was purified using the minelute reaction cleanup kit (Qiagen, 28204)

### NGS sequencing and data processing of the DAP-seq

The DNA-seq libraries were sequenced by Macrogen using Illumina NovaSeq 6000 sequencing plateform. A minimum of 55 million 150 bp paired-end reads were obtained per sample (Table S4). Using CLC Genomic Workbench (Qiagen), reads were cleaned and trimmed of adaptors, ambiguous nucleotides, homopolymers in 3’ and 5’ and for quality (PHRED30) and minimum length. They were then mapped to the *Nicotiana tabacum* TN90 reference genome (GCF_000715135.1_Ntab-TN90_genomic) (Sierro et al., 2014). Peak scores (>2) and p-values (<0.05) were then calculated using the Transcription Factor CHIP-seq tool from CLC Genomic Workbench comparing our samples to the negative control (DNA bound to nlsGFP). Peaks of interests were selected based on their proximity to a gene: 3kb upstream of a START codon and 1kb downstream of a STOP codon.

### Microscopy

Trichomes counting was done by observing leaves under an Observer Z1 epifluorescent microscope (Zeiss) equipped with a GFP, DSRed and DAPI filter. Four circular pieces were cut from 12-15 cm leaves in the center area of the leaf from three independent plants. Samples were fixed in ethanol and stained overnight in Calcolfuor White/Fluorescent Brightener28 (VWR, ICNA0215806701), 1h in 0.01% Fluorol Yellow 088 (Santa Cruz Biotechnology, 81-37-8) and 1h in 0.5% Rodamine B (Merck, 7559) and wash in water three times for 10min.

Zen and ImageJ software were used for post-acquisition image processing. Reporter lines of pGL2 and pAGL66 were studied respectively with the Observer Z1 epifluorescent microscope (Zeiss) with the GFP filter and the Leica Sp8 Stellaris confocal microscope

### Electrophoretic Mobility Shift Assay (EMSA)

Recombinant proteins were expressed in *E. coli* BL21(DE3) cells, which were transformed with expression vectors. Cultures were grown at 30°C to OD600 0.3–0.5, then induced with 0.5 mM isopropyl β-D-1-thiogalactopyranoside (IPTG) for 5 hours. Cells were harvested by centrifugation, frozen, and lysed in a buffer (50 mM Tris-HCl, pH 8, 500 mM NaCl, 30 mM imidazole, 0.5 mM DTT, 1 mM PMSF, lysozyme, and protease inhibitors) by sonication. The lysate was clarified by centrifugation, and the protein was purified using a Ni-NTA column. After washing, proteins were eluted with 300 mM imidazole. Protein concentration was measured using a Qubit fluorometer. For the EMSA assay, complementary oligonucleotides encoding the MEME motif were synthesized, with one oligo tagged with Cystein3 (Table S6). Equimolar concentrations (10 μM) of oligos were mixed in STE buffer (100 mM NaCl, 10 mM Tris-HCl, pH 8, 1 mM EDTA) and annealed using a PCR thermocycler with the following program: 98°C for 3 minutes, 75°C for 1 hour, 65°C for 1 hour, 37°C for 30 minutes, and 25°C for 10 minutes. For binding assays, the double-stranded DNA substrate (1 nM) was incubated with the purified protein in a binding buffer containing 25 mM Tris-HCl (pH 7.5), 9% glycerol, 75 mM NaCl, 5 mM DTT, 5 mM MgCl2, and 1 mg/mL BSA at room temperature for 20 minutes. For negative controls without proteins, SEC buffer (50 mM Tris-HCl, pH 8, 10% glycerol, 50 mM NaCl, 5 mM Na2EDTA) was used. The reaction was separated on a 10% polyacrylamide gel and visualized using a Typhoon scanner (Cytiva).

## References

Aerts N, de Bruijn S, van Mourik H, Angenent GC, van Dijk ADJ (2018) Comparative analysis of binding patterns of MADS-domain proteins in Arabidopsis thaliana. BMC Plant Biology 18: 131

Balkunde R, Pesch M, Hülskamp M (2010) Trichome patterning in Arabidopsis thaliana from genetic to molecular models. Curr Top Dev Biol 91: 299–321

Bartlett A, O’Malley RC, Huang S-sC, Galli M, Nery JR, Gallavotti A, Ecker JR (2017) Mapping genome-wide transcription-factor binding sites using DAP-seq. Nature Protocols 12: 1659–1672

Bereterbide A, Hernould M, Castera S, Mouras A (2001) Inhibition of cell proliferation, cell expansion and differentiation by the Arabidopsis SUPERMAN gene in transgenic tobacco plants. Planta 214: 22–29

Berhin A, de Bellis D, Franke RB, Buono RA, Nowack MK, Nawrath C (2019) The Root Cap Cuticle: A Cell Wall Structure for Seedling Establishment and Lateral Root Formation. Cell 176: 1367–1378.e1368

Bowman JL, Smyth DR, Meyerowitz EM (1989) Genes directing flower development in Arabidopsis. The Plant Cell 1: 37–52

Brinkman EK, Chen T, Amendola M, van Steensel B (2014) Easy quantitative assessment of genome editing by sequence trace decomposition. Nucleic Acids Research 42: e168–e168

Chang J, Wu S, You T, Wang J, Sun B, Xu B, Xu X, Zhang Y, Wu S (2024) Spatiotemporal formation of glands in plants is modulated by MYB-like transcription factors. Nature Communications 15: 2303

Chang J, Yu T, Yang Q, Li C, Xiong C, Gao S, Xie Q, Zheng F, Li H, Tian Z, Yang C, Ye Z (2018) Hair, encoding a single C2H2 zinc-finger protein, regulates multicellular trichome formation in tomato. Plant J 96: 90–102

Choi YE, Lim S, Kim H-J, Han JY, Lee M-H, Yang Y, Kim J-A, Kim Y-S (2012) Tobacco NtLTP1, a glandular-specific lipid transfer protein, is required for lipid secretion from glandular trichomes. The Plant Journal 70: 480–491

Ditta G, Pinyopich A, Robles P, Pelaz S, Yanofsky MF (2004) The SEP4 gene of Arabidopsis thaliana functions in floral organ and meristem identity. Current biology : CB 14: 1935–1940

Dorca-Fornell C, Gregis V, Grandi V, Coupland G, Colombo L, Kater MM (2011) The Arabidopsis SOC1-like genes AGL42, AGL71 and AGL72 promote flowering in the shoot apical and axillary meristems. The Plant Journal 67: 1006–1017

Dreni L, Kater MM (2014) reloaded: evolution of the subfamily genes. New Phytologist 201: 717–732

Ewas M, Gao Y, Wang S, Liu X, Zhang H, Nishawy EME, Ali F, Shahzad R, Ziaf K, Subthain H, Martin C, Luo J (2016) Manipulation of SlMXl for enhanced carotenoids accumulation and drought resistance in tomato. Science Bulletin 61: 1413–1418

Fahn A (2000) Structure and function of secretory cells. In Advances in Botanical Research, Vol 31. Academic Press, pp 37–75

Ferrándiz C, Gu Q, Martienssen R, Yanofsky MF (2000) Redundant regulation of meristem identity and plant architecture by FRUITFULL, APETALA1 and CAULIFLOWER. Development 127: 725–734

Folter Sd, Angenent GC (2006) <em>trans</em> meets <em>cis</em> in MADS science. Trends in Plant Science 11: 224–231

Gao S, Gao Y, Xiong C, Yu G, Chang J, Yang Q, Yang C, Ye Z (2017) The tomato B-type cyclin gene, SlCycB2, plays key roles in reproductive organ development, trichome initiation, terpenoids biosynthesis and Prodenia litura defense. Plant Sci 262: 103–114

Gianfagna TJ, Carter CD, Sacalis JN (1992) Temperature and Photoperiod Influence Trichome Density and Sesquiterpene Content of Lycopersicon hirsutum f. hirsutum 1. Plant Physiology 100: 1403–1405

Glas JJ, Schimmel BC, Alba JM, Escobar-Bravo R, Schuurink RC, Kant MR (2012) Plant glandular trichomes as targets for breeding or engineering of resistance to herbivores. Int J Mol Sci 13: 17077–17103

Gramzow L, Theissen G (2010) A hitchhiker’s guide to the MADS world of plants. Genome Biology 11: 214

Guan Y, Chen S, Chen F, Chen F, Jiang Y (2022) Exploring the Relationship between Trichome and Terpene Chemistry in Chrysanthemum. In Plants, Vol 11

Guo S, Sun B, Looi L-S, Xu Y, Gan E-S, Huang J, Ito T (2015) Co-ordination of Flower Development Through Epigenetic Regulation in Two Model Species: Rice and Arabidopsis. Plant and Cell Physiology 56: 830–842

Han G, Li Y, Yang Z, Wang C, Zhang Y, Wang B (2022) Molecular Mechanisms of Plant Trichome Development. Frontiers in Plant Science 13

Hauser MT (2014) Molecular basis of natural variation and environmental control of trichome patterning. Front Plant Sci 5: 320

Hiratsu K, Ohta M, Matsui K, Ohme-Takagi M (2002) The SUPERMAN protein is an active repressor whose carboxy-terminal repression domain is required for the development of normal flowers. FEBS Letters 514: 351–354

Horsch RB, Fry J, Hoffmann N, Neidermeyer J, Rogers SG, Fraley RT (1989) Leaf disc transformation. In SB Gelvin, RA Schilperoort, DPS Verma, eds, Plant Molecular Biology Manual. Springer Netherlands, Dordrecht, pp 63–71

Huchelmann A, Boutry M, Hachez C (2017) Plant Glandular Trichomes: Natural Cell Factories of High Biotechnological Interest. Plant Physiol 175: 6–22

Jin J, Tian F, Yang D-C, Meng Y-Q, Kong L, Luo J, Gao G (2017) PlantTFDB 4.0: toward a central hub for transcription factors and regulatory interactions in plants. Nucleic Acids Research 45: D1040–D1045

Jin R, Klasfeld S, Zhu Y, Fernandez Garcia M, Xiao J, Han S-K, Konkol A, Wagner D (2021) LEAFY is a pioneer transcription factor and licenses cell reprogramming to floral fate. Nature Communications 12: 626

Karimi M, Inzé D, Depicker A (2002) GATEWAY™ vectors for Agrobacterium-mediated plant transformation. Trends in Plant Science 7: 193–195

Kempin SA, Mandel MA, Yanofsky MF (1993) Conversion of Perianth into Reproductive Organs by Ectopic Expression of the Tobacco Floral Homeotic Gene NAG1. Plant Physiology 103: 1041–1046

Lee J, Lee I (2010) Regulation and function of SOC1, a flowering pathway integrator. Journal of Experimental Botany 61: 2247–2254

Liu X, Bartholomew E, Cai Y, Ren H (2016) Trichome-Related Mutants Provide a New Perspective on Multicellular Trichome Initiation and Development in Cucumber (Cucumis sativus L). Front Plant Sci 7: 1187

Liu Y, Liu D, Khan AR, Liu B, Wu M, Huang L, Wu J, Song G, Ni H, Ying H, Yu H, Gan Y (2018) NbGIS regulates glandular trichome initiation through GA signaling in tobacco. Plant Mol Biol 98: 153–167

Love MI, Huber W, Anders S (2014) Moderated estimation of fold change and dispersion for RNA-seq data with DESeq2. Genome Biology 15: 550

Lu X, Huang L, Scheller HV, Keasling JD (2023) Medicinal terpenoid UDP-glycosyltransferases in plants: recent advances and research strategies. Journal of Experimental Botany 74: 1343–1357

Lv Z, Li J, Qiu S, Qi F, Su H, Bu Q, Jiang R, Tang K, Zhang L, Chen W (2022) The transcription factors TLR1 and TLR2 negatively regulate trichome density and artemisinin levels in Artemisia annua. Journal of Integrative Plant Biology 64: 1212–1228

Ma YJ, Li PT, Sun LM, Zhou H, Zeng RF, Ai XY, Zhang JZ, Hu CG (2020) HD-ZIP I Transcription Factor (PtHB13) Negatively Regulates Citrus Flowering through Binding to FLOWERING LOCUS C Promoter. Plants (Basel) 9

Markus Lange B, Turner GW (2013) Terpenoid biosynthesis in trichomes—current status and future opportunities. Plant Biotechnology Journal 11: 2–22

Matias-Hernandez L, Aguilar-Jaramillo AE, Cigliano RA, Sanseverino W, Pelaz S (2016) Flowering and trichome development share hormonal and transcription factor regulation. J Exp Bot 67: 1209–1219

Meng Q, Gao Y-N, Cheng H, Liu Y, Yuan L-N, Song M-R, Li Y-R, Zhao Z-X, Hou X-F, Tan X-M, Zhang S-Y, Huang X, Ma Y-Y, Xu Z-Q (2025) Molecular mechanism of interaction between SHORT VEGETATIVE PHASE and APETALA1 in Arabidopsis thaliana. Plant Physiology and Biochemistry 220: 109512

Mi H, Muruganujan A, Huang X, Ebert D, Mills C, Guo X, Thomas PD (2019) Protocol Update for large-scale genome and gene function analysis with the PANTHER classification system (v.14.0). Nature Protocols 14: 703–721

Nadakuduti SS, Pollard M, Kosma DK, Allen C, Ohlrogge JB, Barry CS (2012) Pleiotropic Phenotypes of the sticky peel Mutant Provide New Insight into the Role of CUTIN DEFICIENT2 in Epidermal Cell Function in Tomato. Plant Physiology 159: 945–960

Ó’Maoiléidigh DS, Wuest SE, Rae L, Raganelli A, Ryan PT, Kwaśniewska K, Das P, Lohan AJ, Loftus B, Graciet E, Wellmer F (2013) Control of Reproductive Floral Organ Identity Specification in Arabidopsis by the C Function Regulator AGAMOUS The Plant Cell 25: 2482–2503

Pelaz S, Gustafson-Brown C, Kohalmi SE, Crosby WL, Yanofsky MF (2001) APETALA1 and SEPALLATA3 interact to promote flower development. The Plant Journal 26: 385–394

Pnueli L, Hareven D, Broday L, Hurwitz C, Lifschitz E (1994) The TM5 MADS Box Gene Mediates Organ Differentiation in the Three Inner Whorls of Tomato Flowers. The Plant Cell 6: 175–186

Pottier M, Laterre R, Van Wessem A, Ramirez AM, Herman X, Boutry M, Hachez C (2020) Identification of two new trichome-specific promoters of Nicotiana tabacum. Planta 251: 58

Rerie WG, Feldmann KA, Marks MD (1994) The GLABRA2 gene encodes a homeo domain protein required for normal trichome development in Arabidopsis. Genes Dev 8: 1388–1399

Riechmann JL, Meyerowitz EM (1997) MADS domain proteins in plant development. Biol Chem 378: 1079–1101

Schmittgen TD, Livak KJ (2008) Analyzing real-time PCR data by the comparative CT method. Nat. Protocols 3: 1101–1108

Schuurink R, Tissier A (2020) Glandular trichomes: micro-organs with model status? New Phytologist 225: 2251–2266

Sierro N, Battey JND, Ouadi S, Bakaher N, Bovet L, Willig A, Goepfert S, Peitsch MC, Ivanov NV (2014) The tobacco genome sequence and its comparison with those of tomato and potato. Nature Communications 5: 3833

Soltis DE, Chanderbali AS, Kim S, Buzgo M, Soltis PS (2007) The ABC Model and its Applicability to Basal Angiosperms. Annals of Botany 100: 155–163

Szymanski DB, Lloyd AM, Marks MD (2000) Progress in the molecular genetic analysis of trichome initiation and morphogenesis in Arabidopsis. Trends in Plant Science 5: 214–219

Thomas PD, Ebert D, Muruganujan A, Mushayahama T, Albou L-P, Mi H (2022) PANTHER: Making genome-scale phylogenetics accessible to all. Protein Science 31: 8–22

Tissier A, Morgan JA, Dudareva N (2017) Plant Volatiles: Going ‘In’ but not ‘Out’ of Trichome Cavities. Trends in Plant Science 22: 930–938

Vandesompele J, De Preter K, Pattyn F, Poppe B, Van Roy N, De Paepe A, Speleman F (2002) Accurate normalization of real-time quantitative RT-PCR data by geometric averaging of multiple internal control genes. Genome Biol 3: Research0034

Wada T, Tominaga-Wada R (2015) CAPRICE family genes control flowering time through both promoting and repressing CONSTANS and FLOWERING LOCUS T expression. Plant Science 241: 260–265

Wang X, Ye L, Lyu M, Ursache R, Löytynoja A, Mähönen AP (2020) An inducible genome editing system for plants. Nature Plants 6: 766–772

Wang Z, Yan X, Zhang H, Meng Y, Pan Y, Cui H (2021) NtCycB2 negatively regulates tobacco glandular trichome formation, exudate accumulation, and aphid resistance. Plant Molecular Biology

Werker E (2000) Trichome diversity and development. In Advances in Botanical Research, Vol 31. Academic Press, pp 1–35

Wu M, Bian X, Hu S, Huang B, Shen J, Du Y, Wang Y, Xu M, Xu H, Yang M, Wu S (2024) A gradient of the HD-Zip regulator Woolly regulates multicellular trichome morphogenesis in tomato. The Plant Cell 36: 2375–2392

Wu M, Chang J, Han X, Shen J, Yang L, Hu S, Huang B-B, Xu H, Xu M, Wu S, Li P, Hua B, Yang M, Yang Z, Wu S (2023) A HD-ZIP transcription factor specifies fates of multicellular trichomes via dosage-dependent mechanisms in tomato. Developmental Cell 58: 278–288.e275

Wu ML, Cui YC, Ge L, Cui LP, Xu ZC, Zhang HY, Wang ZJ, Zhou D, Wu S, Chen L, Cui H (2020) NbCycB2 represses Nbwo activity via a negative feedback loop in tobacco trichome development. J Exp Bot

Xu J, van Herwijnen ZO, Drager DB, Sui C, Haring MA, Schuurink RC (2018) SlMYC1 Regulates Type VI Glandular Trichome Formation and Terpene Biosynthesis in Tomato Glandular Cells. Plant Cell 30: 2988–3005

Yan T, Chen M, Shen Q, Li L, Fu X, Pan Q, Tang Y, Shi P, Lv Z, Jiang W, Ma Y-n, Hao X, Sun X, Tang K (2017) HOMEODOMAIN PROTEIN 1 is required for jasmonate-mediated glandular trichome initiation in Artemisia annua. New Phytologist 213: 1145–1155

Yang C, Gao Y, Gao S, Yu G, Xiong C, Chang J, Li H, Ye Z (2015) Transcriptome profile analysis of cell proliferation molecular processes during multicellular trichome formation induced by tomato Wov gene in tobacco. BMC Genomics 16: 868

Yang C, Li H, Zhang J, Wang T, Ye Z (2011) Fine-mapping of the woolly gene controlling multicellular trichome formation and embryonic development in tomato. Theor Appl Genet 123: 625–633

Yang C, Ye Z (2013) Trichomes as models for studying plant cell differentiation. Cell Mol Life Sci 70: 1937–1948

Yang S, Miao H, Zhang S, Cheng Z, Zhou J, Dong S, Wehner TC, Gu X (2011) Genetic analysis and mapping of gl-2 gene in cucumber (Cucumis sativus L.). Acta Horticulturae Sinica 38: 1685–1692

Zhang B, Wang L, Zeng L, Zhang C, Ma H (2015) Arabidopsis TOE proteins convey a photoperiodic signal to antagonize CONSTANS and regulate flowering time. Genes & Development 29: 975–987

Zheng F, Cui L, Li C, Xie Q, Ai G, Wang J, Yu H, Wang T, Zhang J, Ye Z, Yang C (2021) Hair (H) interacts with SlZFP8-like to regulate the initiation and elongation of trichomes by modulating SlZFP6 expression in tomato. Journal of Experimental Botany

