## Supplemental Figures for "From Genes to Glands: Unraveling the Pivotal Influence of NtAGL66, an AGAMOUS-like Transcription Factor, on Glandular Trichome Development in *Nicotiana tabacum*"

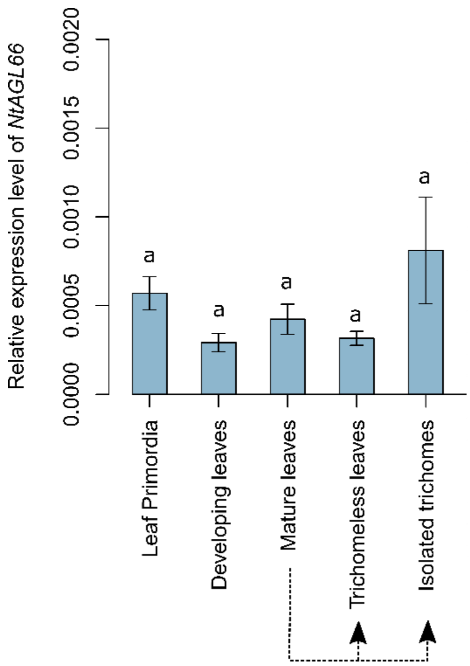


**Figure S1: Expression level of *NtAGL66* in different tissues.** RT-qPCR on different tissues of WT plants. The tissues selected for analysis encompass various stages of leaf development and trichome presence, including leaf primordia (early leaf development stage), developing (i.e. still in the elongation phase) leaves, mature leaves, isolated trichomes (both glandular and potentially non-glandular), and leaves without trichomes (shaved leaves). *NtAGL66* display a very low expression level compared to internal control genes in all the tissues we investigated (the expression level is actually very close to background noise This might be due to the fact that this gene could be expressed in a highly specific manner, limited to a small number of cells within the plant. Gene expression levels are expressed relative to that corresponding to the geometric mean of three different internal controls (*NtEF1a, NtTAC9, NtUBC2*). Results are displayed as mean ± SD, n=3-4. Significant differences were determined by ANOVA. Each letter represents a significant variation between a tissue ($p < 0.05$).

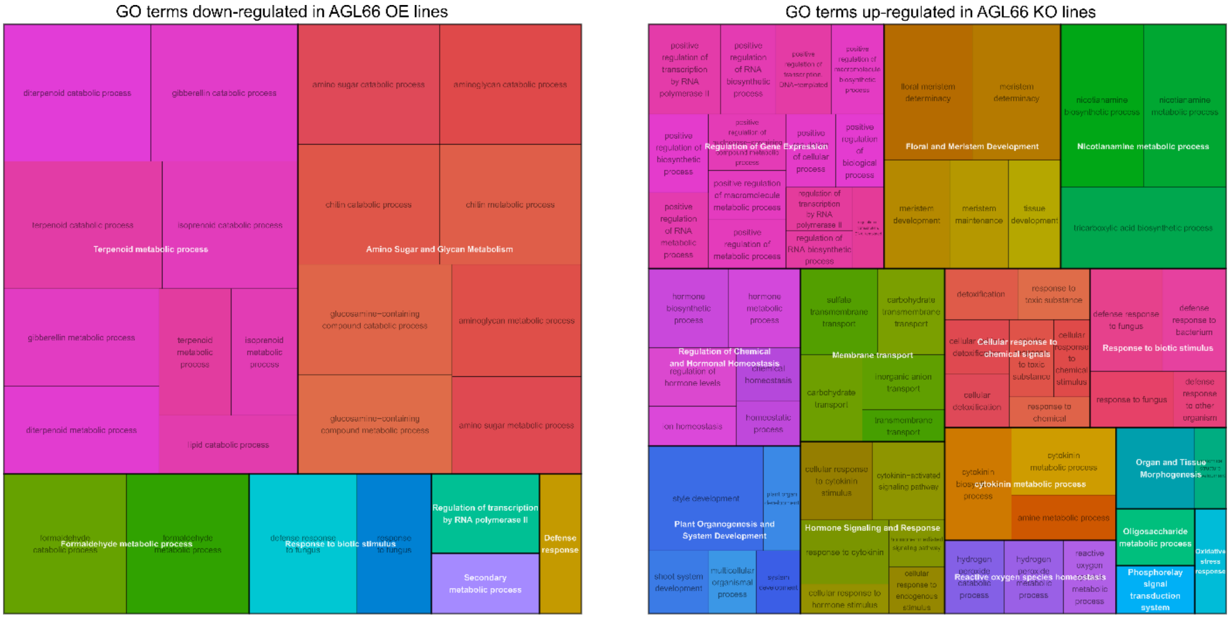


**Figure S2: Groups of Go terms inversely correlated with NtAGL66 expression.** Overview of the enriched categories using Rrvigo (Sayols, 2023), a tool to help visualize enriched GO terms.

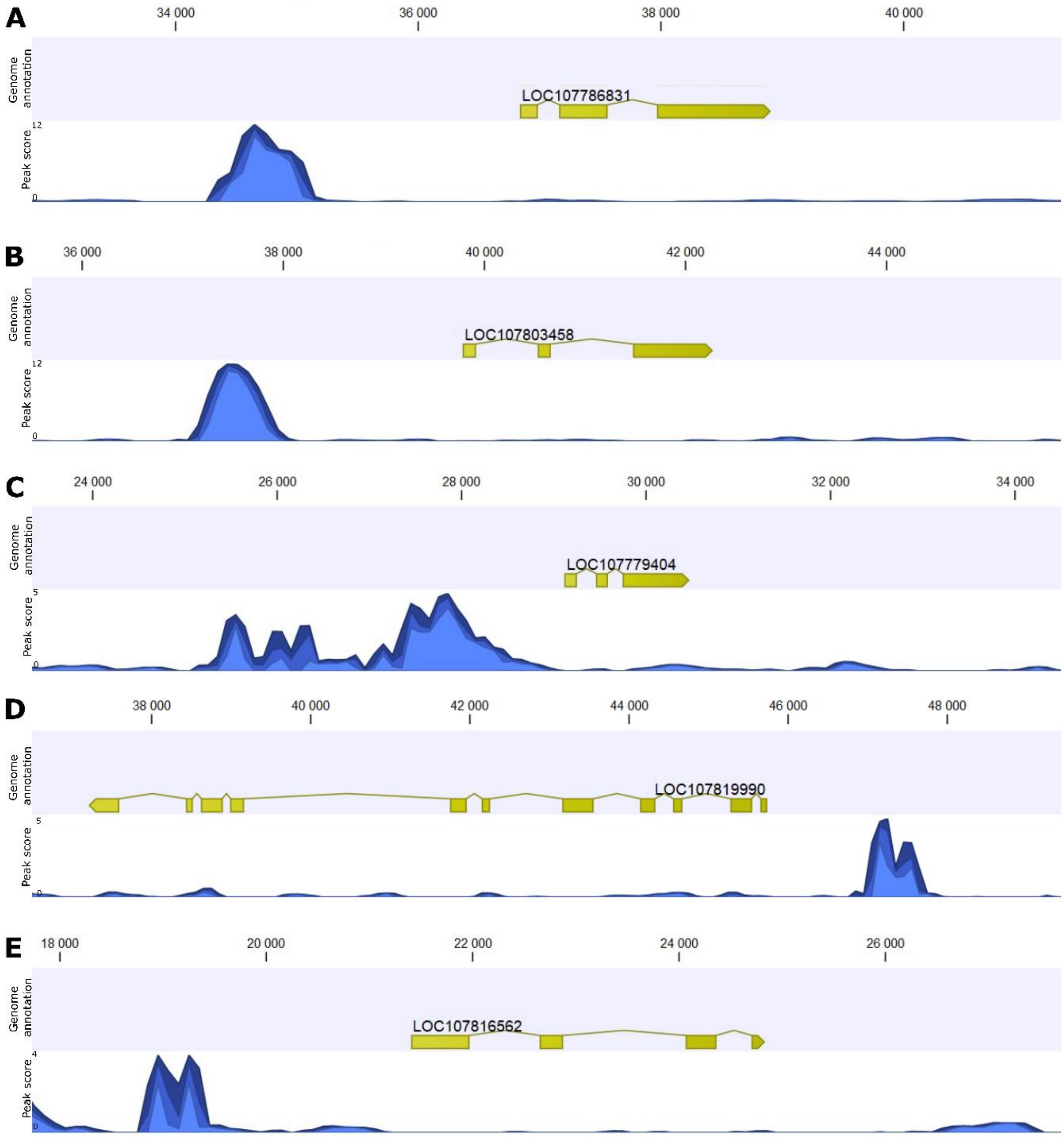


**Figure S3: NtAGL66 binding sites of the TOP5 transcription factors bound in their promoter.** Illustration of the binding region of (A) LOC107786831, NtJACKDAW; (B) LOC107803458, NtMYB6; (C) LOC77799404, RAX3-like;(D) LOC107819990, NtGLABRA2; (E)LOC107816562, NtCONSTANS-like 10. In each of these panels, the upper track presents the gene annotation and its position on the contig while the lower track presents the DAP-seq peak shape score and the localization of the peak. The analysis and this figure were done on CLC Genomics.
